# ARID4 Modulates Stomatal ABA Signaling and Stress-Responsive Gene Expression to Confer Drought Tolerance in Arabidopsis

**DOI:** 10.64898/2026.06.16.732254

**Authors:** Zhiqing Zhang, Zhengyu Shao, Yilin Yang, LiaoLiao Ye, Yajie Liu, Yiji Xia, Tao Qin, Liming Xiong

## Abstract

To identify novel regulators of plant drought tolerance, we performed a forward genetic screen using thermal infrared imaging to detect alterations in leaf temperature, a proxy for transpiration. This screen identified AT-rich Interacting Domain Protein 4 (ARID4), a protein involved in chromatin remodeling, as a modulator of transpiration. *arid4* loss-of-function mutants exhibited cooler leaves, increased water loss from detached leaves, and heightened susceptibility to drought relative to wild-type plants. Stomata of *arid4* mutants were hyposensitive to abscisic acid (ABA), displaying reduced ABA-induced reactive oxygen species (ROS) production and impaired stomatal closure. Transcriptome profiling under drought revealed extensive misregulation of genes involved in stress-responses, redox homeostasis, iron acquisition, and photosynthesis. These findings indicate that ARID4 modulates the expression of key regulatory genes to maintain redox homeostasis and photosynthetic efficiency, thereby conferring drought stress tolerance.

## Introduction

Soil water deficit (drought) is a common environmental stress often encountered by land plants. Plants have evolved an array of strategies to mitigate the negative impacts of drought. Among these strategies, transpiration control is a rapid and crucial response for plants to survive drought episodes. Transpiration control is achieved through the closure of stomatal pores, which are formed by pairs of guard cells. Although guard cell signaling is extensively studied, our knowledge of transpiration regulation remains limited due to technical challenges and the inherent complexity of the process.

Stomatal movement is mainly mediated by ion and solute transport across guard cell membranes and the resulting water movement that changes the turgidity of the cells (Ward et al. 2009). Transporters involved in the process can be quickly regulated through posttranslational modifications, particularly phosphorylation. The stress phytohormone abscisic acid (ABA), which accumulates under water deficit or osmotic stress conditions plays a critical role in modulating these posttranslational modifications. Once ABA is perceived by its PYR/PYL/RCAR receptors, protein kinases such as SnRK2s are activated to phosphorylate membrane transporters and regulate ion and solute movement (Cutler et al. 2010; Chen et al. 2020). Consequently, posttranslational modifications of transporters are considered the norm in regulating stomatal movement (Schroeder et al. 2013) whereas transcriptional regulation has received little attention. Nonetheless, over the past decades, transcription factors have also been found to participate in stomatal regulation, although the mechanisms are still under investigation (Liang et al. 2005; Cominelli et al. 2005). The discovery of transcription factors in stomatal regulation suggests that proteins involved in chromatin remodeling and gene activation may also play roles in stomatal movement as well. However, progress in this area remains limited and innovative approaches aimed at revealing transpiration regulation mechanisms are needed.

Since transpiration decreases leaf temperatures, genetic screening of leaf temperatures using infrared cameras would potentially allow direct access to genes critical for transpiration regulation. However, not many genes were previously identified using this method (Mustilli et al. 2002; Liang et al. 2010; Merlot et al. 2007; Dong et al. 2018), although the method was successful in identifying key determinants of stomatal CO_2_ responses (Hashimoto et al. 2006; Negi et al. 2008; Negi et al. 2013; Takahashi et al. 2025). By improving the genetic screening method, we obtained new mutants with altered leaf temperatures. The first locus we characterized is *LOT1* (*lo*wer *t*emperature *1*) that encodes a protein involved in regulating the subcellular distribution and stability of ABA-signaling components (Qin et al. 2019). The second locus identified, *LOT2* (re-named as *ARID4*, see below), which encodes an AT-rich interaction domain (ARID) protein, will be the focus of this study.

AT-rich interaction domain (ARID) proteins are DNA-binding proteins found in all eukaryotes with a preference for binding AT-rich DNA sequences. The ARID domain does not share sequence homology to other known DNA-binding domains but was demonstrated to bind DNA, often in a less sequence-specific manner (Iwahara et al. 2002). Proteins in this group are essential components of SWI/SNF chromatin-remodeling complexes and are often found mutated in cancers (Centore et al. 2020). Mutations in these genes also affect development, particularly that of reproductive systems (Lahoud et al. 2001; Wu et al. 2013). These ARID protein-containing complexes have a modular organization whereby different compositions of the complex give rise to different functionalities, e.g., regulating the expression of different sets of genes (Raab et al. 2015). The ARID protein is the defining component of these chromatin-remodeling complexes and also facilitates ATPase subunit binding (Mashtalir et al. 2018). Upon recruitment by sequence-specific transcription factors, the ARID protein-containing chromatin remodeling complexes often target to specific regulatory regions to modulate spatiotemporal gene expression (Ho et al. 2019).

The Arabidopsis genome encodes 14 ARID proteins. In addition to the ARID domain, their C-termini often harbor an additional domain such as the PHD, ELM2, HMG, or Hsp20 domains (Zhu et al. 2008). Similar to their functions in animals, the few studies on functions of ARID proteins in plants also point to their involvement in development and association with chromatin. For instance, ARID1, which associates with a histone deacetylase complex is involved in sperm formation (Zheng et al. 2014). The ARID domain in ARID5 recognizes the histone H3K4me3 mark and is required for ARID5 to interact with chromatin in regulating floral transition (Tan et al. 2020). Other ARID proteins are involved in shoot meristem development (Xu et al. 2015), heterochromatin formation (Tan et al. 2018), pollen development (Xia et al. 2014; Zheng et al. 2014), and nodulation (Zhu et al. 2008). Although how ARID proteins regulate these developmental processes is unknown, gene regulation is likely the major mechanism both in animals and in plants. Indeed, ARID proteins bind DNA and interact with transcription factors. For example, the ARID-containing chromatin remodeling BAF components in animals can directly interact with nuclear hormone receptors (Ho et al. 2019) to direct gene expression, while OsARID3 interacts with the CONSTANS-like protein Ghd2 to affect development and senescence (Liu et al. 2016).

In this study, we found that the *ARID4* locus is essential for maintaining plant drought stress tolerance through the control of stomatal movement. The *arid4* mutant had a cooler leaf temperature, and its stomata were less responsive to ABA in eliciting reactive oxygen species (ROS) production and stomatal closure. RNA-seq analysis of wild-type and mutant plants subjected to drought stress treatments indicates that the expression of many stress-responsive and redox-related genes was misregulated in the mutant. Furthermore, certain chloroplast genes involved in photosynthesis were significantly downregulated, whereas certain iron uptake and transport genes were upregulated under drought stress but not under normal conditions. Peculiarly, the ARID4 protein was also partially localized in chloroplast under osmotic stress conditions, ARID4 appears to regulate gene expression and stomatal movement using a pathway that is not entirely dependent on ABA. Together, our results suggest that ARID4 has unique functions in plant drought stress tolerance, likely through controlling ROS homeostasis and maintaining photosynthetic efficiency under drought stress.

## Results

### Isolation of *arid4* as a transpiration-defective mutant

Using infrared cameras to monitor leaf temperatures, we screened both EMS-mutagenized and T-DNA mutagenized populations (Alonso et al. 2003) of *Arabidopsis thaliana* (ecotype Columbia-0) for leaf temperature alterations. One T-DNA insertional mutant (SALK_007400) was found to have markedly lower leaf temperatures compared with the wild type. We initially named the mutant *lot2* (*lo*w *t*emperature *2*). The T-DNA was inserted in an exon of a gene encoding an ARID/BRIGHT DNA-binding domain-containing protein, which was later annotated as *ARID4* in the database. We therefore renamed the mutant *arid4* for consistency. In multiple experiments, the temperatures of *arid4* leaves were approximately 0.7 to 1.0 °C lower than those of the wild type (based on multipoint measurements across the canopy) (Figure 1A). Consistent with the lower temperatures, detached leaves of *arid4* had higher water loss (transpiration) rates than the wild type (Figure 1B).

**Figure 1.**
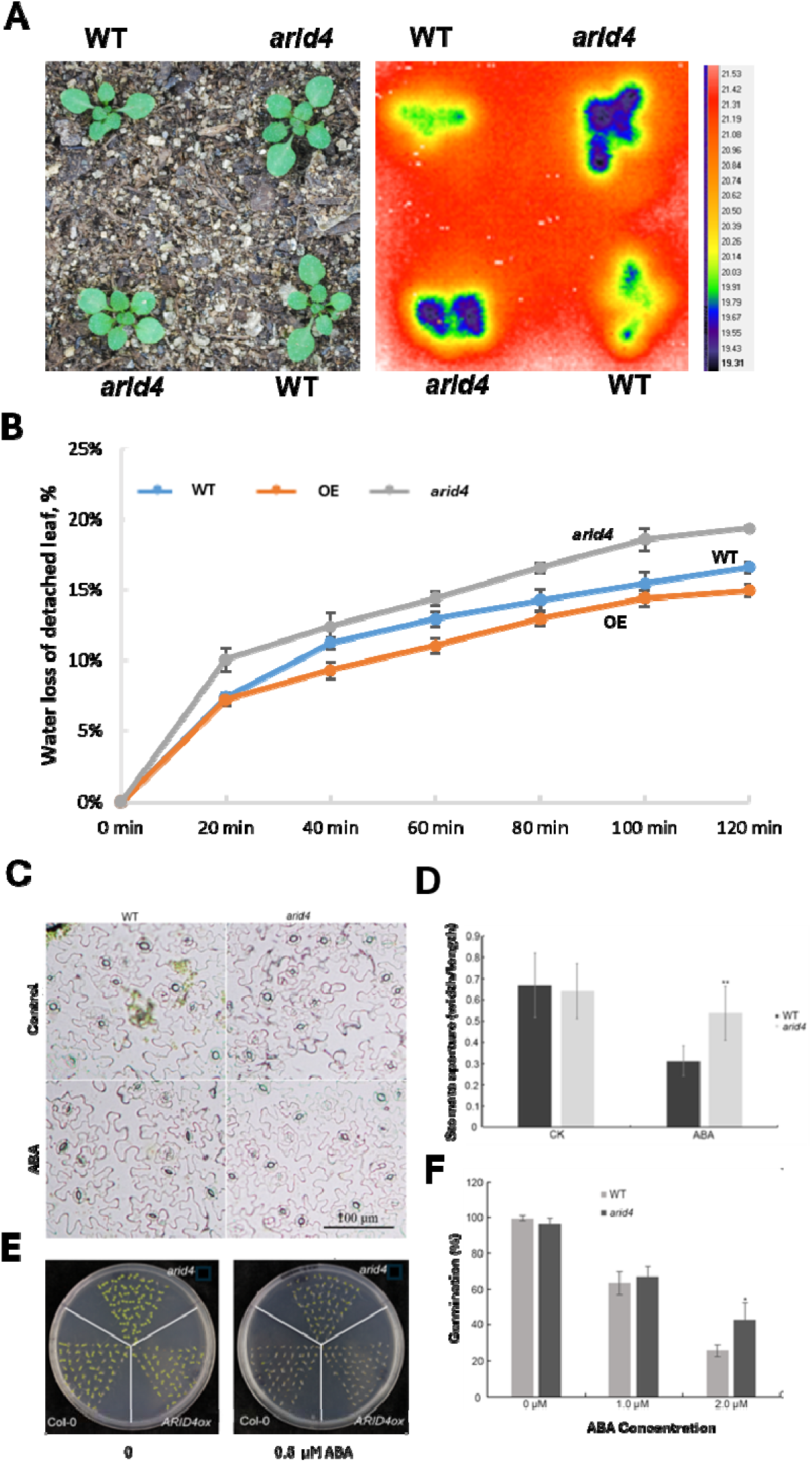
Isolation of the *arid4* mutant and characterization of its ABA responses. **A**. Morphology of the wild type (WT) and *arid4* seedlings and their infrared images. The pseudocolor scale indicates leaf surface temperature, ranging from 19.3°C (blue) to 21.5°C (red). Representative images from one of three independent experiments are shown. **B**. Water loss rate of detached leaves. Water loss was measured as the percentage of initial fresh weight lost over time. **C**. Representative images of stomata from WT and *arid4* plants after a 2-h treatment with 20 µM ABA. Scale bar = 100 µm. **D**. Quantification of stomatal aperture. Epidermal strips with open stomata were incubated in buffer with or without 20 µM ABA for 2 h. Aperture is expressed as the width-to-length ratio. Data are from approximately 500 stomata per treatment. Asterisks indicate significant differences from the WT under the same condition (Student’s *t*-test, \*\**P* < 0.001). **E**. ABA sensitivity of seed germination. Seeds were sown on ½ MS medium containing 1% sucrose with or without 0.5 µM ABA. Pictures were taken four days after stratification. **F**. Quantification of germination rate. Germination was scored as radicle emergence. Data are presented as mean ± SD from three independent biological replicates. Asterisks indicate a significant difference from the WT under the same condition (Student’s *t*-test, \**P* < 0.01). WT, wild type Col-0; *arid4*/*arid*-T, *arid4* mutant; OE/*ARID4ox*, overexpressing *ARID4* in the Col-0 background.

To investigate the cause of its higher transpiration rate, we examined stomatal density of *arid4* mutant leaves but found no difference from that of the wild type (Supplementary Figure S1). Thus, the higher transpiration rate in *arid4* should result from deregulated stomatal movement. It is known that drought stress increases cellular ABA levels, which triggers the closure of stomata. To determine stomatal responses to ABA, epidermal strips of leaves were treated with ABA, and the stomatal apertures were measured. While wild type stomata responded to ABA by significantly decreasing their aperture, *arid4* mutant stomata were much less responsive, and the aperture did not decrease significantly (Figures 1C-D).

Consistent with reduced stomatal sensitivity to ABA, the germination of *arid4* seeds was also much less inhibited by ABA (Figure 1E). Seed germination rates with 1.0 or 2.0 µM ABA treatment were higher than those of the wild type, and the difference was significant for the higher ABA concentration (Figure 2F). Interestingly, while short term treatment of ABA similarly inhibited the root elongation of both the wild type and *arid4* (Supplementary Figure S2), osmotic stress significantly inhibited lateral root growth of *arid4* mutants (Figures 2A-B). Since both ABA and osmotic stress inhibit lateral root growth (Xiong et al. 2006), the distinct response of *arid4* mutant roots to osmotic stress suggest that ARID4 may regulate responses to osmotic stress differently from its role in ABA signaling.

**Figure 2.**
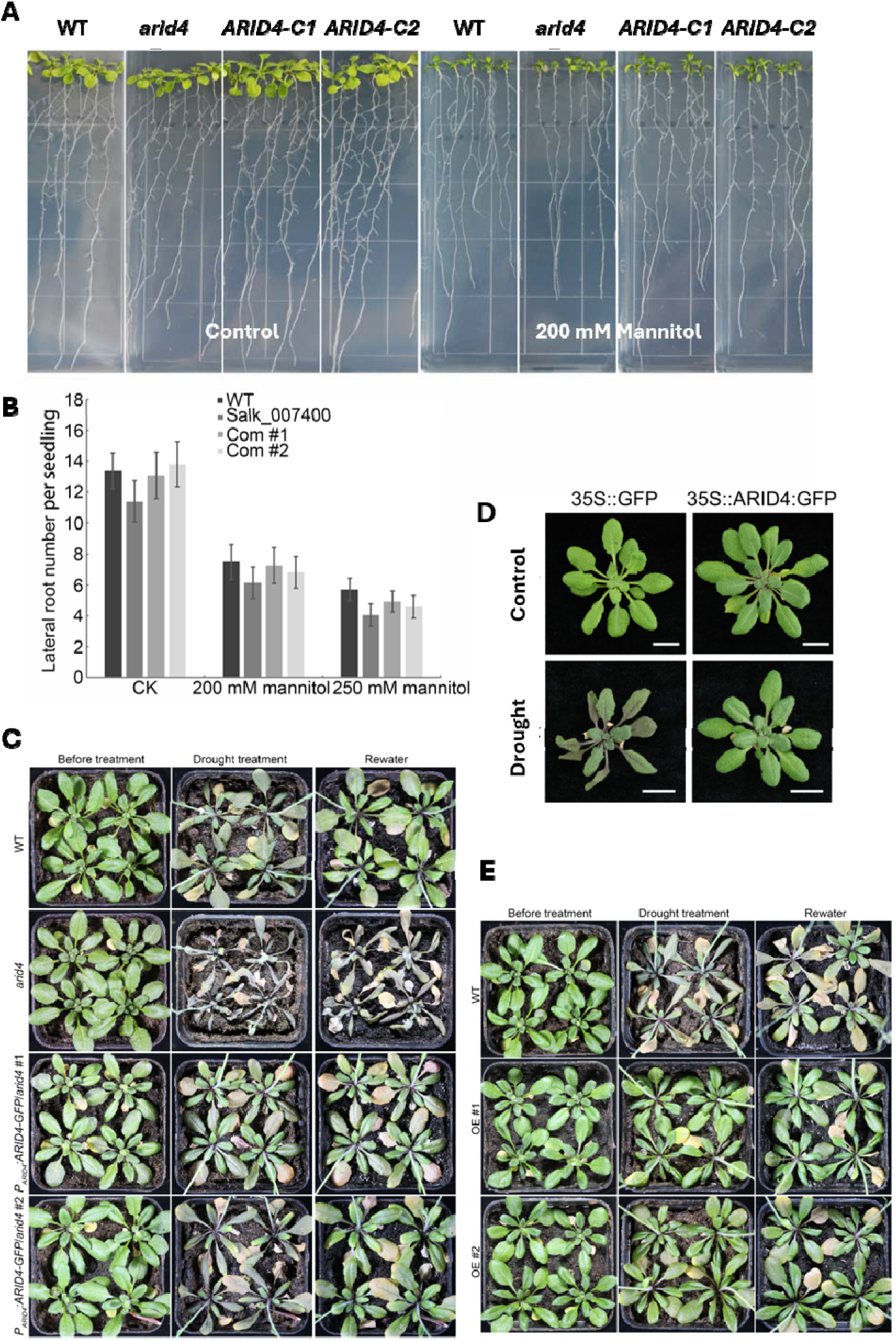
Response of the *arid4* mutant to osmotic stress and drought. **A**. Root growth response to osmotic stress generated by 200 mM mannitol. **B**. Quantitation of lateral root numbers shown in **A**. Error bars are SD. **C**. The *arid4* mutant was drought-sensitive and the wild type ARID4 complemented its drought-sensitive phenotypes. Three-week-old seedlings were stopped watering for 17 days and then re-watered. Pictures were taken before and after the drought treatment and 2 days after recovery. **D-E**. Drought tolerance of transgenic plants overexpressing *ARID4*. Soil-grown seedlings after withholding water for 11 days (**D**) or 17 days, and 2 days after rewatering (**E**). WT, wild type Col-0; P*_ARID4_*.*ARID4-GFP*, wild type *ARID4* fused with *GFP* under the control of native *ARID4* promoter in *arid4* mutant background; OE/OE#1 or #2 over-expressing *ARID4* in Col-0 background, line #1 or #2; ARID4-C1/Com #1 or #2, *ARID4* complementation line #1 or

### The *arid4* mutant is sensitive to drought stress

The data presented above show that *arid4* mutant stomata are less responsive to ABA and that mutant leaves lost water more rapidly than the wild type. We therefore tested the drought sensitivity of soil-grown *arid4* plants compared to the wild type. Plants were allowed to grow for three weeks with full irrigation and then subjected to water withholding for 17 days. At this point, all plants had wilted to varying degrees. The plants were then re-watered, and photographs were taken 2 days later (Figure 2C). Whereas the wild-type plants fully recovered from the drought stress, most of the *arid4* mutant plants failed to recover.

To confirm that the phenotypes observed in *arid4* mutants were caused by the loss of the wild type *ARID4* gene, we generated complementation lines. A construct was made with the wild-type *ARID4* gene fused with *GFP* under the control of the native *ARID4* promoter. The construct was introduced into the *arid4* mutant and homozygous transgenic seedlings were obtained through selection and confirmation. These transgenic plants were subjected to the drought stress treatment described above and were found to recover from the stress similarly to wild-type plants (Figure 2C). Furthermore, their lateral roots showed similar response to mannitol as the wild type (Figures 2A-2B). These data indicate that loss of the wild type *ARID4* gene was responsible for these phenotypic defects seen in the *arid4* mutant.

### Overexpression of *ARID4* Enhances Drought Resistance

We also generated transgenic plants expressing the wild type *ARID4* gene fused with the GFP reporter gene under the control of the constitutive CaMV 35S promoter. The construct was transferred into the wild-type background, and homozygous transgenic plants were obtained. When these plants were subjected to mild stress (water withheld for 10 days), the leaves of control transgenic plants (expressing an empty GFP vector) turn purple, yet the *ARID4* overexpressing plants remained green (Figure 2D). When three-week-old seedlings were subjected to more severe drought (water withheld for 17 days), the wild-type plants showed clear leaf withering, whereas the leaves of the *ARID4* overexpressing plants remained turgid. Two days after rewatering, although the wild type plants recovered, some of their leaves had senesced, and increased anthocyanin accumulation was observed. These stress-related phenotypes were less evident in the overexpressing plants (Figure 2E). These data demonstrate that overexpressing *ARID4* could enhance plant drought resistance.

### Expression pattern of *ARID4* and subcellular localization of the ARID4 protein

To investigate the expression pattern of the *ARID4* gene, stable transgenic plants expressing the *GUS* reporter gene driven by the native *ARID4* promoter were generated. GUS signals were observed at relatively low levels in roots, leaves, flowers, and siliques (Figure 3). However, strong signals were observed in guard cells (Figure 3). To determine protein localization, a construct with an ARID4-GFP fusion under the control of the native *ARID4* promoter was introduced into *arid4* mutant plants. The fusion protein was consistently localized to the nucleus in primary roots in all transgenic plants (Figure 4A). ARID-GFP signals were also readily observed in the primordial cells of lateral roots (Figure 4A) and in the nuclei of guard cells (Figure 4B), whereas the expression level was low or absent in other cell types (e.g., non-meristematic root cells and mesophyll cells). Interestingly, ARID4-GFP signals in root cells were found in the nucleoplasm but were excluded from the nucleolus in non-dividing cells (Figure 4A).

**Figure 3.**
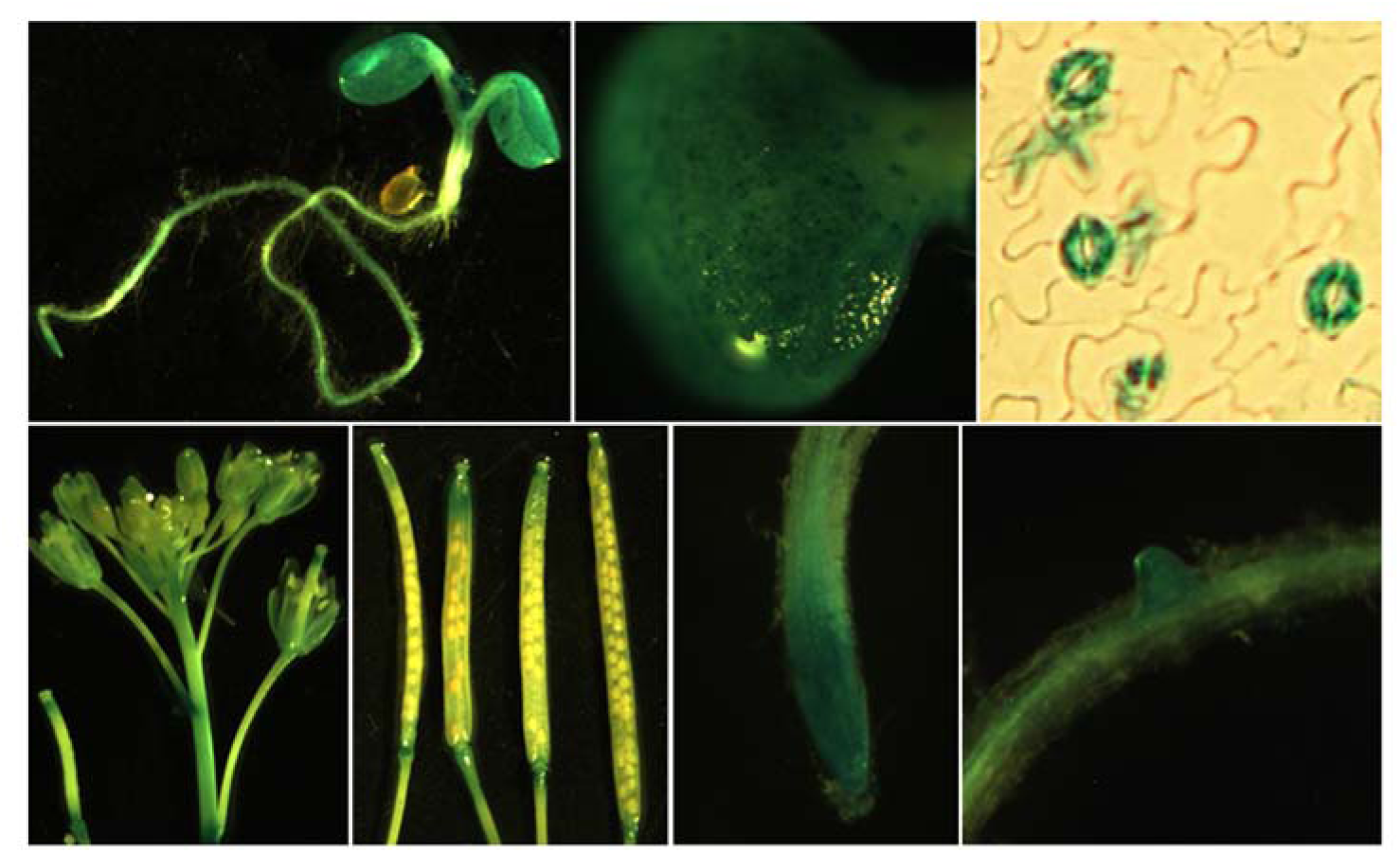
Expression pattern of *pARID4-GUS*. Histochemical GUS staining of transgenic *Arabidopsis* plants expressing the *pARID4::GUS* reporter construct. Representative images show GUS activity (blue) in young seedlings, guard cells, flowers, siliques, primary and lateral roots.

**Figure 4.**
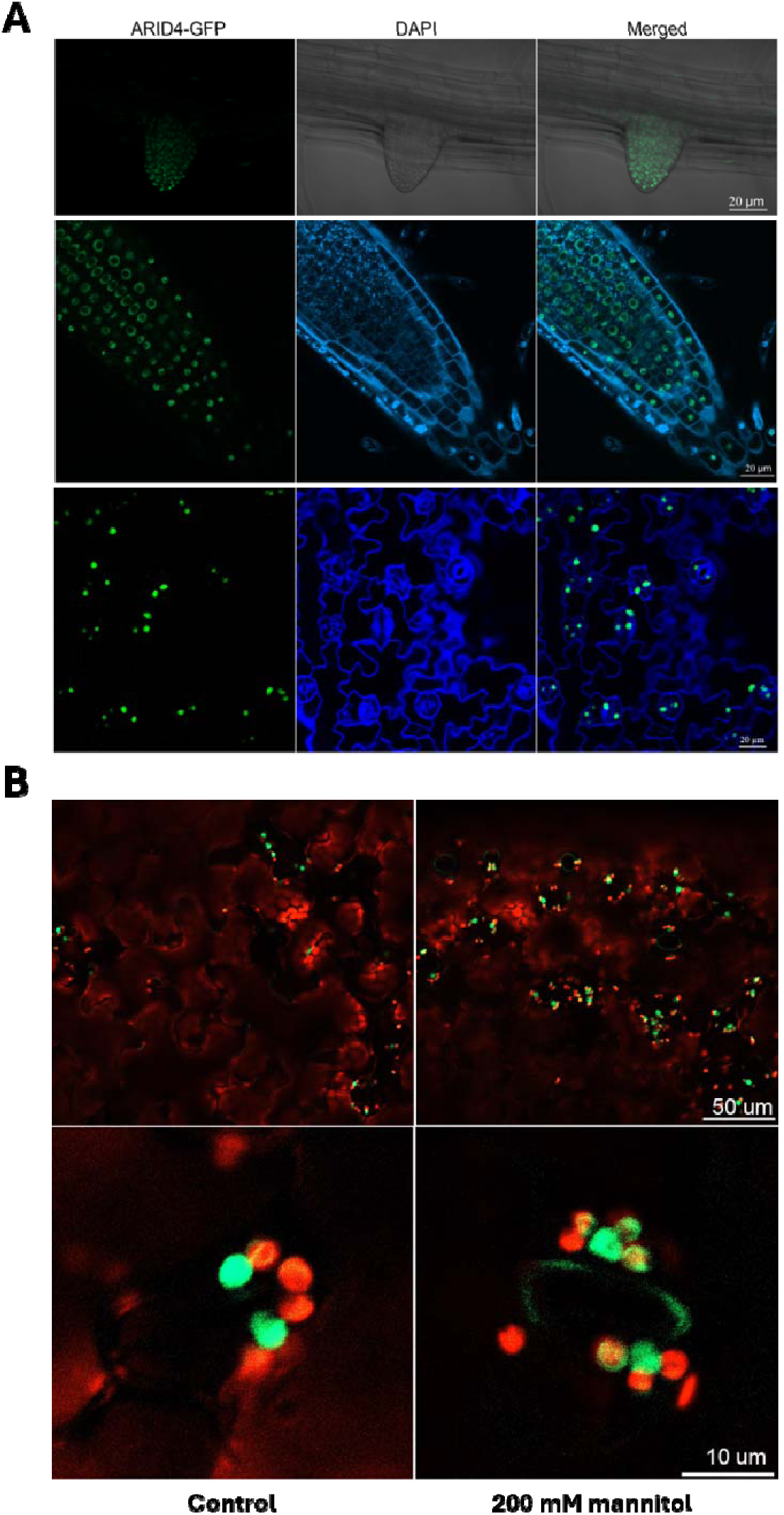
Subcellular localization of ARID4-GFP. Confocal microscopy of transgenic Arabidopsis expressing *pARID4::ARID4-GFP* in the *arid4* mutant background. **A**. GFP signals were detected in nuclei of lateral (upper panel) and primary (middle panel) root cells, and in guard cells (lower panel). The green channel shows ARID4-GFP fluorescence, the blue signals show DAPI staining, and “Merge” shows the combined signals. **B**. Under normal (mock-treated) conditions, ARID4-GFP is localized to the nucleus in guard cells (left panels). After treatment with 200 mM mannitol to induce osmotic stress, ARID4-GFP was observed in the nucleus and also partly co-localized with chlorophyll autofluorescence, indicating partial localization to chloroplasts. Scale bars = 20 µm in **A**, and 10 or 50 µm in **B** as shown.

We also investigated whether the localization of the ARID4 protein was affected by ABA or stress treatments. While we did not observe clear changes in ARID4-GFP localization under ABA treatments, it was noted that osmotic stress, generated by 200 mM mannitol treatment, induced partial localization of the GFP signal to chloroplasts in guard cells (Figure 4B). This localization was observed in independent assays and suggests that the ARID4 protein, despite being a primarily nuclear protein, may have additional functions in chloroplasts under osmotic stress conditions.

### *arid4* guard cells are less responsive to ABA in inducing ROS production

ABA is known to induce the production of H_2_O_2_ and trigger stomatal closure. Since *arid4* stomata are less responsive to ABA in inducing stomatal closure (Figures 1C-1D), we examined whether *arid4* is defective in ABA-induced ROS production. Leaves of *arid4* were incubated in 10 µM ABA, and ROS levels were visualized by staining with H_2_DCFDA (2′,7′-dichlorodihydrofluorescein diacetate) and observed under a confocal microscope. As shown in Figure 5, under control conditions, weak ROS staining was similarly observed in leaves of the wild type, *arid4* and *ARID4*-overexpressing plants. After ABA treatment, higher levels of fluorescence were readily observed compared with the untreated samples. Although there was some interference from chlorophyll autofluorescence, ABA-induced ROS could be observed in the cytoplasm and near the plasma membrane. The level of ROS production was lowest in the *arid4* mutant, highest in the *ARID4*-overexpressing plants, and intermediate in the wild type. The level of ABA-induced fluorescence correlates with the degree of stomata closure and water loss rates of detached leaves (Figures 1B and 5), indicating that *arid4* had reduced response to ABA in generating ROS. This defect may contribute to its reduced stomatal closure in response to ABA.

**Figure 5.**
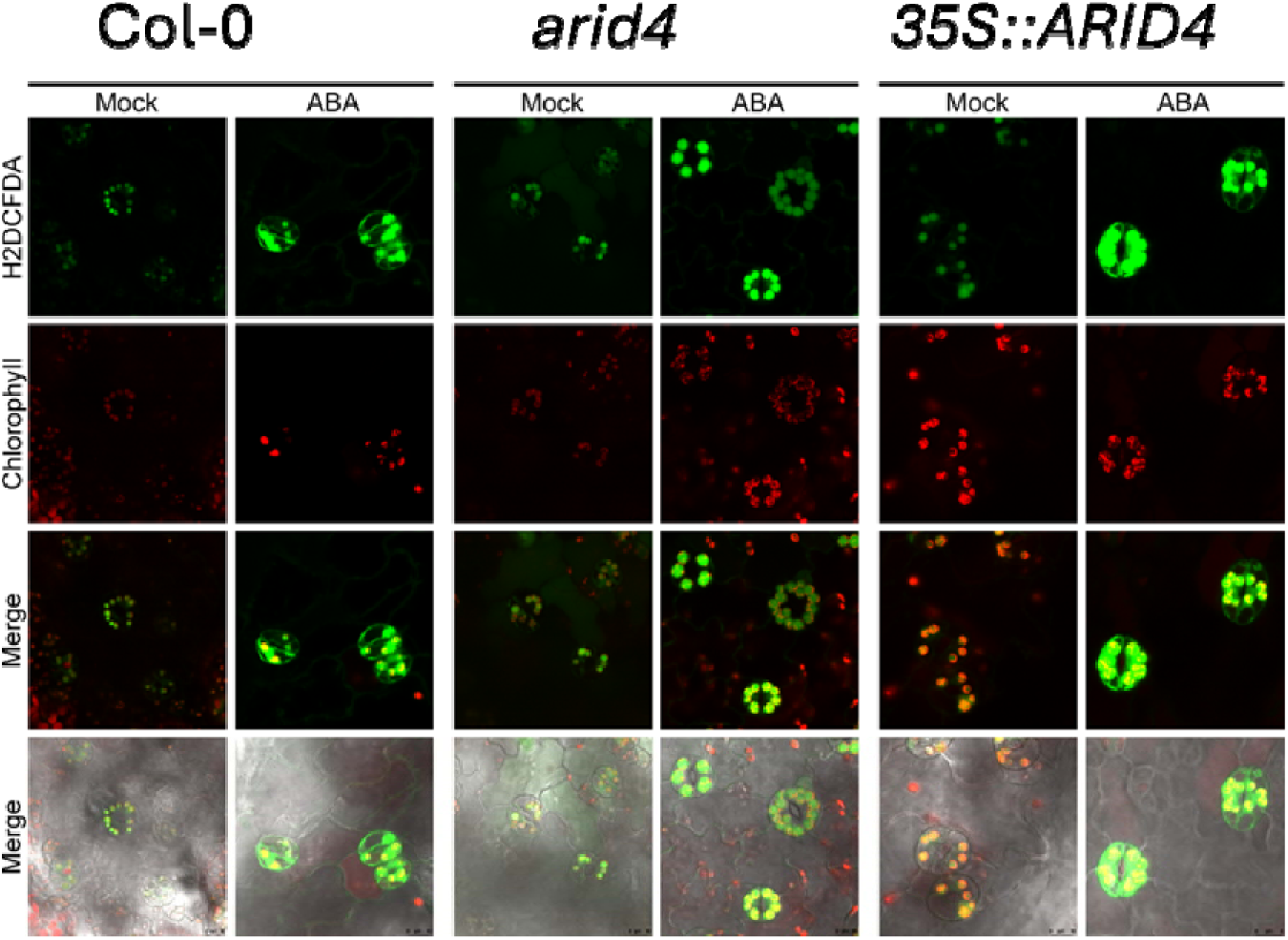
The *arid4* mutant is impaired in ABA-induced ROS production in guard cells. Representative confocal images showing reactive oxygen species (ROS) production in the guard cells of wild-type (Col-0), *arid4,* and *ARID4* overexpressing (Col-0 background) (*35S::ARID4*) plants. Epidermal peels were treated with a control buffer (Mock) or 10 µM ABA for 1 hour, then stained with 10 µM H_2_DCFDA for 20 min to detect ROS (visualized as green fluorescence).

### Transcriptome analysis of drought-stressed *arid4* mutants

In animals, ARID homologs are DNA-binding proteins that affect gene expression through either chromatin remodeling or direct DNA-binding abilities. In Arabidopsis, ARID4 has been found in histone deacetylation complexes that participate in heterochromatin silencing and deacetylation of other loci (Tan et al. 2018; Feng et al. 2021). While several other *ARID* genes in plants were investigated for their role in development, their potential regulation of gene expression under abiotic stresses is less explored. Since *arid4* mutants did not show clear developmental defects under normal conditions yet are clearly involved in drought stress responses, we investigated the potential impacts of ARID4 on gene expression under drought conditions.

To this end, leaves of *arid4* and wild-type plants were harvested and allowed to lose 40% of their fresh weight and then incubated under light for three hours in a sealed plastic bag to prevent further water loss before being harvested and frozen in liquid nitrogen for RNA extraction. RNA-seq analysis showed that under control (non-drought) conditions, 285 genes were differentially expressed in *arid4* compared with the wild type. Among them, 114 genes were significantly upregulated and 171 genes down regulated in the mutant. Under drought stress conditions, 325 genes were differentially regulated in *arid4*, with 118 genes were downregulated and 207 genes up regulated compared with the wild type (Figures 6A and 6C). Functional group analysis of these genes revealed that they belong to a wide range of biological processes, such as “response to oxidative stress”, “oxidation reduction”, “response to abiotic stimulus”, and “defense response”.

**Figure 6.**
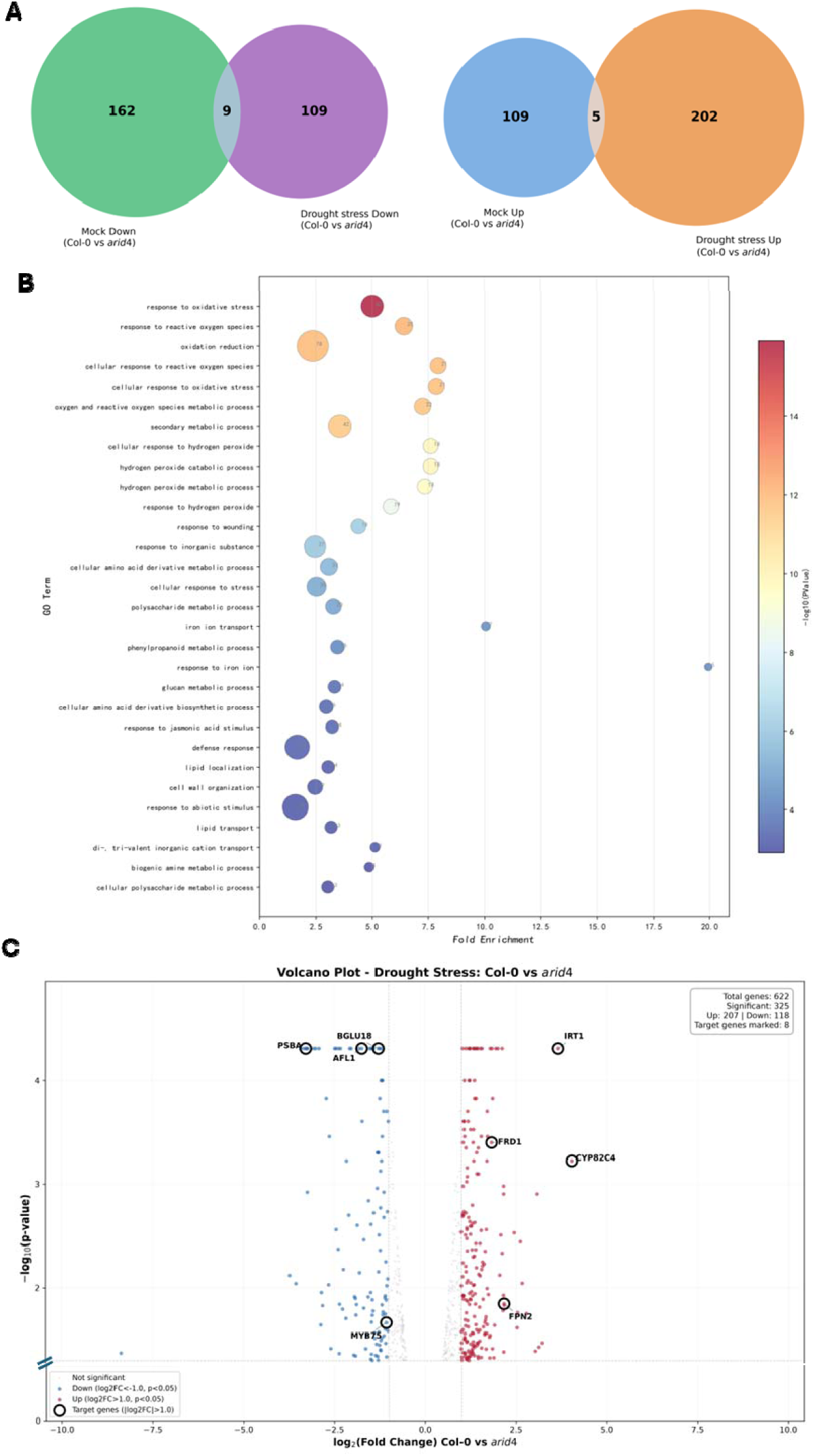
Differentially expressed genes in the *arid4* mutant compared with the wild type under drought stress treatment. RNA-seq was performed with seedlings treated with or without drought stress. Differentially expressed genes (DEGs) were defined as |log_2_ (fold change)| ≥ 1 and adjusted *p*-value < 0.05. **A**. Total number of downregulated (left) or upregulated (right) DEGs identified between the wild type and *arid4* under control or drought treatment. **B**. Gene Ontology (GO) enrichment of DEGs in the *arid4* relative to the wild type under drought treatment. The *x*-axis shows the mean log_2_ (fold change) of genes associated with each term, the *y*-axis lists significant GO terms (grouped by biological process), bubble size is proportional to the number of DEGs in each group, and color intensity reflects the –log_10_ (*p*-value). **C**. Volcano plot comparing transcript abundance between wild type (Col-0) and *arid4* under drought treatment. Each dot represents a gene; x-axis: log_2_ (fold change); y-axis: –log_10_ (*p*-value). Upregulated and downregulated DEGs are shown in red and blue, respectively. Genes whose expression level were confirmed with RT-qPCR are labeled with gene identifiers.

Intriguingly, several genes involved in iron uptake and transport were significantly upregulated and highly enriched in the mutant under drought (Figure 6B). Furthermore, certain photosynthesis genes, particularly those in photosystem II (PSII) were down regulated in *arid4* compared with the wild type under drought stress conditions.

### Validation of Gene Expression by RT-qPCR

To validate the transcriptomic data, we chose several differentially expressed genes to perform quantitative real-time PCR (RT-qPCR) assays. Seedlings grown on MS agar medium were harvested and dehydrated to induce a 40% loss of fresh weight, then incubated for three hours under high humidity conditions. A separate set of seedlings were sprayed with 100 µM ABA. RT-qPCR analysis confirmed the RNA-seq results for the selected genes. For instance, several known stress-responsive genes, including the drought tolerance gene *AFL1* (AT3g28270) (Kumar et al. 2015), *DREB2B* (Liu et al. 1998), *WHY1*, *DRIR1* (Qin et al. 2017), and *RAB18*, were confirmed to be down-regulated in *arid4* under drought or ABA treatment. In contrast, iron uptake or transport-related genes, such as *IRT1*, *FRO2*, *CYP82C*, and *FPN2,* which were upregulated in the RNA-seq data, were also confirmed by RT-qPCR to be upregulated under drought stress. The chloroplast gene *PsbA* was also confirmed to be downregulated in *arid4* under drought but not under ABA treatment (Figure 7). These data validate the accuracy of our RNA-seq analysis, and the expression patterns of several of these key genes are indicated in the volcano plot (Figure 6C).

**Figure 7.**
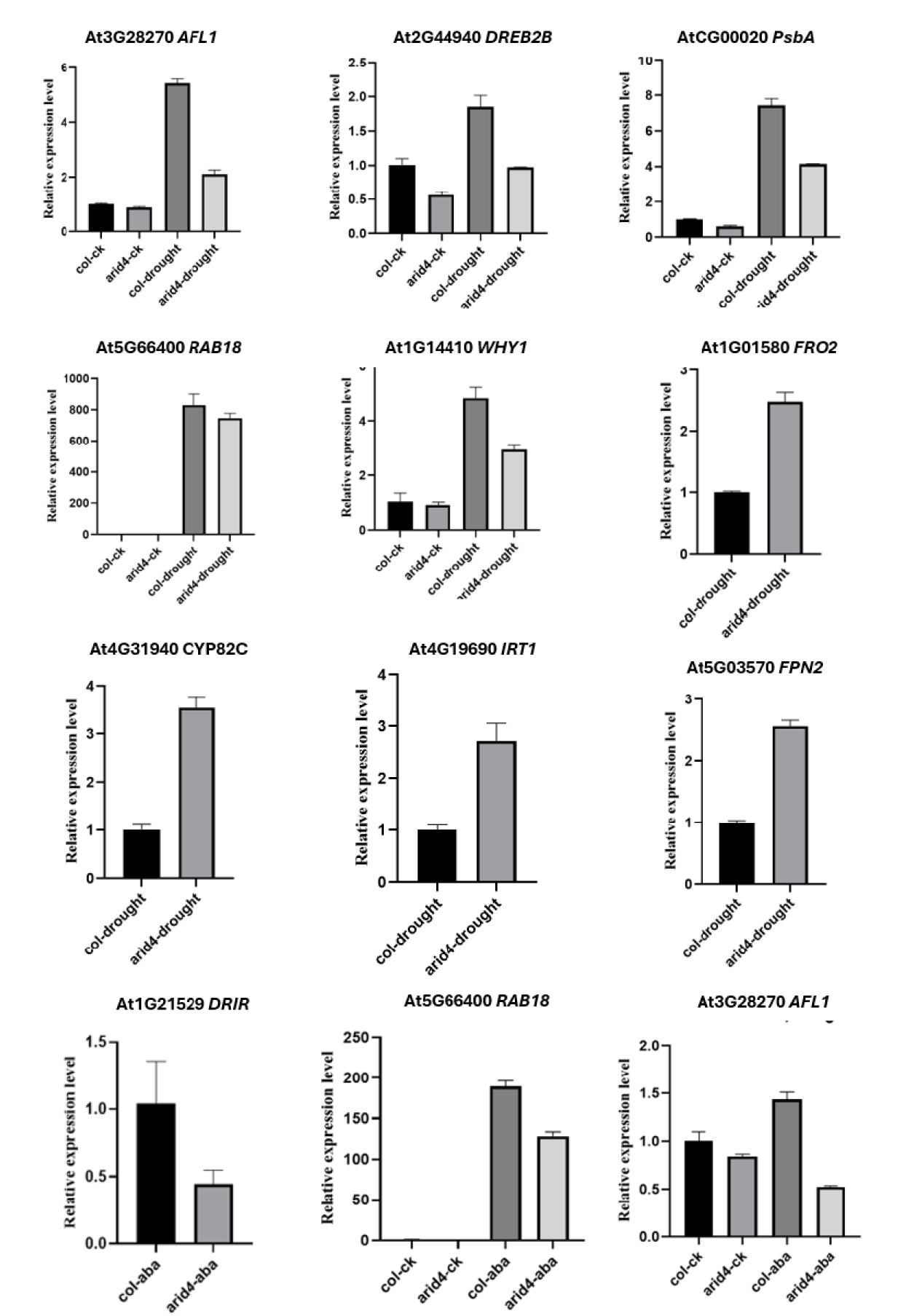
RT-qPCR validation of RNA-seq data for selected differentially expressed genes in *arid4*. Relative expression levels of representative upregulated and downregulated genes in wild-type (Col) and *arid4* mutant plants. RNA was extracted from leaves of plants under control conditions or after dehydration treatment (40% fresh weight loss). Expression levels were normalized to the *ACTIN2* internal control and are shown relative to the WT under control conditions (set to 1). Data are presented as mean ± SD from three biological replicates.

### ARID4 May Help Maintain Photosynthetic Efficiency Under Drought Stress

As a protein involved in chromatin-remodeling, it is not surprising that ARID4 may be involved in multiple biological processes. Yet, the *arid4* mutant does not exhibit obvious phenotypes under normal conditions. Transcriptomic analysis under drought stress revealed altered gene expression in multiple processes, though their functional consequences are not immediately clear. For instance, 78 genes in the ‘oxidation reduction’ category were differentially expressed in *arid4* under drought stress (Figure 6B), suggesting potentially altered sensitivity to oxidative stress in the mutant. Similarly, the higher expression of iron uptake and transport genes suggests a potential disruption of iron homeostasis. While the molecular mechanisms for these transcriptomic changes and their potential impacts are still being investigated, we examined the functional consequences of the reduced expression of photosynthetic genes using chlorophyll fluorescence imaging. Under mild drought conditions (11 days water withholding, when leaves remained turgid), the mutant exhibited chlorophyll fluorescence parameters, such as *Fv/Fm* (maximum quantum yield of PSII) and *NPQ* (non-photochemical quenching), that were mostly indistinguishable from those of the wild type, suggesting normal PSII function under these conditions. Nonetheless, *Y(II)* (the effective quantum yield of PSII) showed a notable reduction in *arid4* (Figure 8), indicating that the capacity for CO_2_ fixation and electron flow was beginning to decline in the mutant even at this early stage of drought stress.

**Figure 8.**
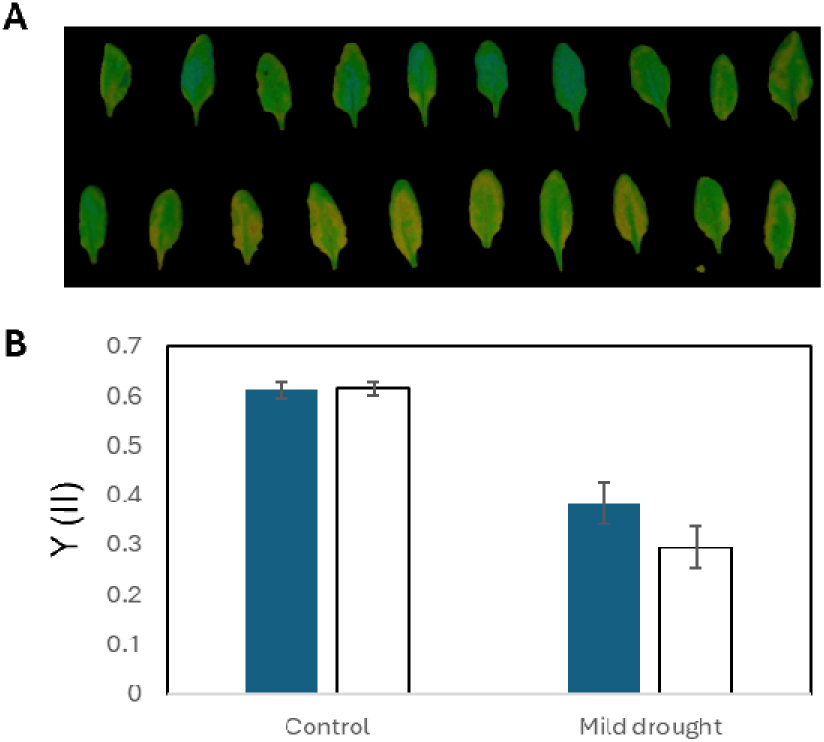
PSII photochemical efficiency in wild-type and the *arid4* mutant. **A**. False-color image of effective quantum yield [*Y(II)*] under growth light in control (well-watered) and drought conditions. The drought treatment was mild, and no clear wilt phenotype was seen among the plant. **B**. Quantification of *Y(II)*. Data are means and standard deviations.

## Discussion

Our genetic screen identified ARID4 as an important regulatory component in controlling leaf transpiration and drought stress tolerance. The stomata of the *arid4* mutant are less responsive to ABA, perhaps partly because of the mutant’s reduced production of ABA-induced H_2_O_2_, which triggers stomatal closure (Zhang et al. 2001; Pei et al. 2000). Nonetheless, the altered transcriptome in *arid4* under drought stress also suggests that ARID4 reprograms gene expression to mitigate the impact of drought. This likely involve the remodeling of chromatin at important drought tolerance loci that control ROS production and detoxification, stress gene activation, iron homeostasis and photosynthesis.

Among the differentially regulated genes, over 50 belong to the GO category ‘response to abiotic stimulus’. Significantly down-regulated genes included well-known players involved in drought stress tolerance, such as *DREB2B* (Liu et al. 1998), *AFL1*(Kumar et al. 2015), and the long non-coding RNA *DRIR* (Qin et al. 2017).

Intriguingly, the *DRIR* transcript, which was nearly undetectable in the *arid4* mutant under drought stress treatment, was recently found to interact with two chloroplast targeted proteins, CP29A and CP29B, and to affect freezing tolerance (Ye et al. 2026).

Abiotic stress often causes the accumulation of reactive oxygen species (ROS). In drought-stressed *arid4* mutants, many genes involved in redox processes were differentially expressed. These are various peroxidase, oxidoreductase, dehydrogenase genes, and cytochrome P450 genes. Some of these genes are also involved in secondary metabolic processes, forming another large GO category. While some of these secondary metabolites contribute to drought stress tolerance, others, such as coumarins, facilitate the acquisition of iron from the soil (Fourcroy et al. 2014; Schmid et al. 2014).

The significant up regulation of iron deficiency-induced genes in the *arid4* mutant is intriguing. These genes are mainly involved in coumarin synthesis and transport, as well as iron reduction and transport, all of which help plants take up iron under iron-deficient conditions. For instance, CYP82C4 hydroxylates fraxetin to generate sideretin (Murgia et al. 2011; Wang et al. 2025; Rajniak et al. 2018). These secreted coumarins help mobilize insoluble iron in the rhizosphere for root uptake (Robe et al. 2021). While it is possible that ARID4-mediated histone deacetylation normally keep the iron uptake system in check to prevent iron overload, it is also plausible that *arid4* mutant plants are in a state of iron deficiency, even though no visible symptoms were observed. We are currently investigating the mechanisms and significance of these gene expression changes.

ARID4, a DNA binding protein found in chromatin-remodeling complexes (Tan et al. 2018), is primarily localized to the nucleus. Interestingly, we observed its partial relocation to chloroplasts under osmotic stress conditions (Figure 4B). Several transcription factors are known to be dual-localized in the nucleus and chloroplasts (Qin et al. 2025; Xin et al. 2021; Lin et al. 2020; Krause et al. 2005) and affect chloroplast gene expression. In *arid4* mutants, a number of genes encoding components of photosystem PSII and PSI, the electron transport chain, and the ATP synthesis apparatus are downregulated under drought stress, yet no chlorophyll biosynthesis genes are affected. This is distinct from H4 acetyltransferase NuA4 (Nucleosome acetyltransferase of H4) complex, which controls chlorophyll biosynthesis (Barrero-Gil et al. 2022; Bieluszewski et al. 2022; Zhou et al. 2022). Thus, ARID4’s role may be to help maintain photosynthetic function or efficiency during stress, rather than regulating chlorophyll biosynthesis. Drought stress is known to result in the disassembly of PSII supercomplexes and the degradation of PSII core. Our initial assays showed that with the onset of drought stress, the *arid4* mutant plants exhibited a reduced effective quantum yield of PSII (*Y(II)*), a parameter known to be very sensitive to drought and actually was suggested to be used for early detection of drought stress (Hu et al. 2023). Our ongoing efforts are focused on investigating the role of ARID4 in maintaining photosynthesis under more severe and long-term drought conditions.

## Materials and Methods

### Plant Materials and Growth Conditions

The *Arabidopsis thaliana* ecotype Columbio-0 (Col-0), referred to as the wild type, was the genetic background for all mutants and transgenics used in this study. The mutant screen using infrared imaging was performed as described previously (Qin et al. 2019). The *arid4* mutant (SALK_007400) was backcrossed once with Col-0 and homozygosity was confirmed by genotyping. For plate-based assays, seeds were surface-sterilized with a solution of 50% commercial bleach containing 0.01% Triton X-100 for 5 min, then rinsed at least five times with sterile water. The sterilized seeds were sown on 1/2 Murashige and Skoog (MS) medium plates supplemented with 1% sucrose. After a 3-day stratification at 4°C, the plates were placed in a growth room under a 16-hlight/8-h-dark cycle at 21°C for germination and growth. For soil-based assays, seedlings were transferred to soil and grown under the same conditions.

### Drought Treatment for RNA Analyses

Four-week-old soil-grown plants were used for drought treatment. Leaves were harvested and incubated under light until they lost approximately 40% of their initial fresh weight. The dehydrated leaf samples were then placed in weigh boats and sealed in plastic zipper bags containing water-soaked tissue paper to maintain nearly saturated humidity. The samples were kept physically separated from the wet tissue paper to prevent direct rehydration while minimizing further water loss. After incubation under light for an additional 3 h, the samples were immediately collected for RNA extraction.

### RT-qPCR Analysis

Total RNA was extracted using TRIzol reagent (Invitrogen) following the manufacturer’s instructions. RNA concentration and purity were determined spectrophotometrically (Thermo Fisher Scientific). The isolated RNA was reverse transcribed using a FastKing RT kit with gDNase (TIANGEN). RT-qPCR was performed using SYBR qPCR Master Mix (Vazyme) on a StepOnePlus real-time PCR system (Applied Biosystems). Three experimental replicates were performed per reaction. *ACTIN2* was used as an internal reference gene. The gene-specific primers used for RT-qPCR are listed in Supplemental Table S1.

### Library Construction, Sequencing, and Transcriptome Analysis

Using TRIzol reagent (Invitrogen), total RNAs were extracted from drought-treated and control seedlings. RNA-seq libraries were constructed using the Illumina Whole Transcriptome Analysis Kit following the standard protocol and were sequenced on the Illumina HiSeq platform to generate paired-end reads of 101 bp. TopHat (Trapnell et al. 2012) was used to align mRNA-seq reads against the Arabidopsis genome (TAIR10) with default parameters. Genes with differential expression were identified by using the Cufflinks software package (Trapnell et al. 2012).

### ROS Staining

Abscisic acid (ABA)-induced ROS production in guard cells was detected using the fluorescent probe H_2_DCFDA (Beyotime, S0034S). Specifically, 7-day-old Arabidopsis seedlings were treated with 10 μM ABA in MES buffer (pH 5.8) for 60 min; control seedlings were treated with the same buffer without ABA. After the treatment, seedlings were immersed in a staining buffer containing 10 µM H_2_DCFDA in PBS for 20 min, followed by washing with PBS for 1 min. The epidermal layer was carefully stripped and mounted for immediate observation using a confocal microscope.

### Confocal Microscopy

The H_2_DCFDA fluorescence was visualized using a Leica DMI8 confocal microscope. The excitation and emission wavelengths were 488 nm and 510–540 nm for H_2_DCFDA, and 638 nm and 640–690 nm for chlorophyll fluorescence, respectively.

### Chlorophyll Fluorescence Measurements

Two-week-old soil-grown seedlings were subjected to drought by withholding water. Eleven days later, chlorophyll fluorescence imaging was performed using a HEXAGON-IMAGING-PAM fluorometer (Heinz Walz GmbH, Effeltrich, Germany) under light conditions. Fluorescence parameters were averaged across ten fully expanded rosette leaves. The effective quantum yield of PSII [*Y(II)*] was calculated as (Fm′– F)/Fm′.

## Acknowledgements

This work was supported by a grant from the Research Grants Council (RGC) of the Hong Kong Special Administrative Region, China General Research Fund #12101521 (to L.X).

## Author contribution

L.X. and T.Q. planned and designed the research. L.X., T.Q., and Y.X supervised the work. T.Q., Z.Z, Z.S., Y.Y., L.Y., and Y.L performed experiments. L.X. and T.Q. prepared the manuscript draft. All authors read and approved its content.

## Supplementary Materials

**Figure S1.**
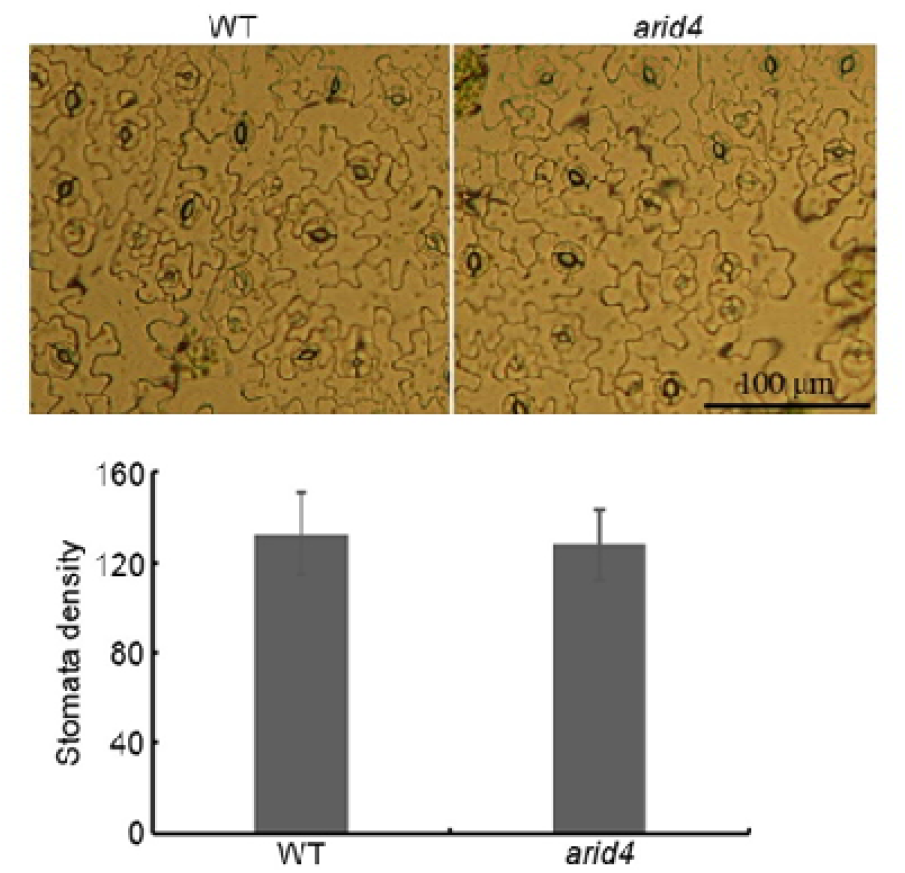
Stomatal density of the wild type and *arid4* mutant.

**Figure S2.**
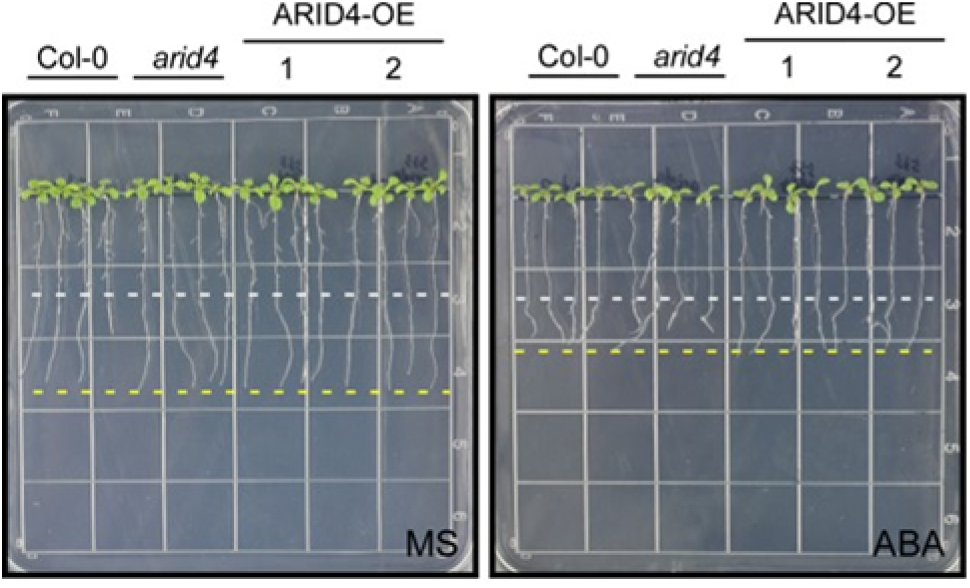
Root growth response to 10 µM ABA in *arid4* and *ARID4* overexpressing plants. ARID4-OE (Line #1 and #2), transgenic plants overexpressing *ARID4* in the Col-0 background.

**Table S1.**
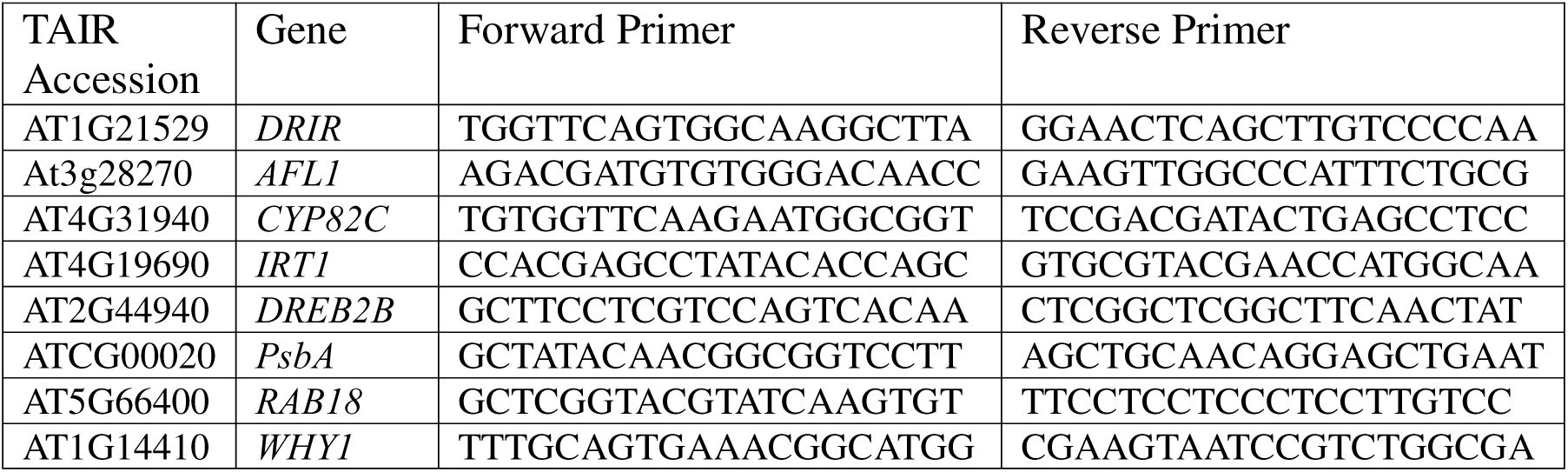
List of primers used in RT-qPCR assays.

